# Inhibitory feedback enables predictive learning of multiple sequences in neural networks

**DOI:** 10.1101/2023.08.26.554928

**Authors:** Matteo Saponati, Martin Vinck

## Abstract

Anticipating future events is a key computational task for neuronal networks. Experimental evidence suggests that reliable temporal sequences in neural activity play a functional role in the association and anticipation of events in time. However, how neurons can differentiate and anticipate multiple spike sequences remains largely unknown. We implement a learning rule based on predictive processing, where neurons exclusively fire for the initial, unpredictable inputs in a spiking sequence, leading to an efficient representation with reduced post-synaptic firing. Combining this mechanism with inhibitory feedback leads to sparse firing in the network, enabling neurons to selectively anticipate different sequences in the input. We demonstrate that intermediate levels of inhibition are optimal to decorrelate neuronal activity and to enable the prediction of future inputs. Notably, each sequence is independently encoded in the sparse, anticipatory firing of the network. Overall, our results demonstrate that the interplay of self-supervised predictive learning rules and inhibitory feedback enables fast and efficient classification of different input sequences.

## Introduction

The prediction of future events is a key mechanism for the organization of behaviour [1], and is arguably supported by anticipatory neuronal firing observed in various brain regions [2, 3, 4, 5, 6]. In order to fire ahead of predictable events, neurons must learn to credit synapses that carry information about future inputs, and downregulated the ones that are predictable. However, this prediction process is difficult when different spiking patterns influence the activity of the same neuronal populations. For instance, synaptic inputs carrying relevant information for predicting one stimulus may also be predictable by other synaptic inputs during another stimulus. To address this issue, neurons must accomplish two tasks: (a) learn predictive relationships in the input spikes patterns, and adjust synaptic weights accordingly; (b) discriminate different components of input statistics and participate in the prediction of specific input patterns. Currently, it remains unclear how neurons can show anticipatory firing that is specific to the particular temporal relationships in the input.

Theoretical works indicate that spike-timing-dependent plasticity (STDP) leads, in some cases, to a reduction in the post-synaptic spike latency when spike patterns are systematically repeated [7, 8]. Thereby, neurons can fire ahead of future sensory stimuli and can give rise to predictions at the network level [9], suggesting a mechanism for anticipatory firing in the brain [2]. Another recent work addressed the problem directly, hypothesizing that neurons adjust their synaptic weights proportionally to the predictability of pre-synaptic inputs [10]. This plasticity mechanism enables neurons to anticipate and signal future events and unifies several STDP phenomena from the perspective of predictive processes. However, this mechanism alone cannot generate anticipatory firing when the temporal relationships between spikes are ambiguous, e.g. when the same input predicts other inputs in one spike sequence but is fully predictable during another sequence. Introducing competition among neurons offers a solution to this issue, as it allows neurons to become selective for specific sequences. Winner-take-all-like models capture the fundamental principle of competition, where single neurons encode specific inputs while other neurons remain silent [11]. A possible competitive mechanism is a recurrent inhibition, which exerts inhibitory control over neurons receiving the same input [12, 13]. Previous studies suggest that local inhibitory interactions, observed in the neocortex [14] and hippocampus [15], are essential for decorrelated neuronal firing and independent representation of input components [16]. However, how the interplay between inhibitory mechanisms and local learning rules can support the prediction of future inputs is still not well understood.

In this work, we propose that the combination of a predictive learning rule (PLR) as in [10] with a competitive mechanism at the network level (inhibitory feedback; IF) can lead to selective anticipation of input patterns. We demonstrate that neurons in the network suppress the firing of the entire network while adjusting their synaptic weights to anticipate predictable inputs. This inhibitory mechanism allows neurons to become selective for specific sequences in the input patterns and to show anticipatory firing for those sequences. Moreover, the identity of different input sequences can be read out from a few early spikes in the network activity, enabling fast and energy-efficient anticipation of future events.

## Results

### Inhibitory feedback enables selective anticipation of spike sequences in neural networks

The present work builds further on a predictive learning rule from a previous work [10] (see Methods for a detailed description of the learning rule). The learning rule potentiates synaptic weights associated with predictive spikes, that is, inputs that anticipate other pre-synaptic inputs, while it suppressed inputs that are anticipated by other pre-synaptic spikes. For example, we explored the scenario where a neuron endowed with the predictive learning rule received a temporal sequence composed of N pre-synaptic neurons firing sequentially with fixed delays (Figure 1a). In this case, the pre-synaptic inputs occurring earlier in the sequence should be potentiated as they anticipate the subsequent inputs. We trained the neuron by repeating the input sequence for 750 epochs. As anticipated, during training, the neuron progressively potentiated the inputs occurring earlier in the sequence. Eventually, it assigned the highest credit to the first inputs in the sequence and fired upon their occurrence (Figure 1b). Notably, our results remained consistent regardless of the size of the pre-synaptic population (Figure S1).

**Figure 1:**
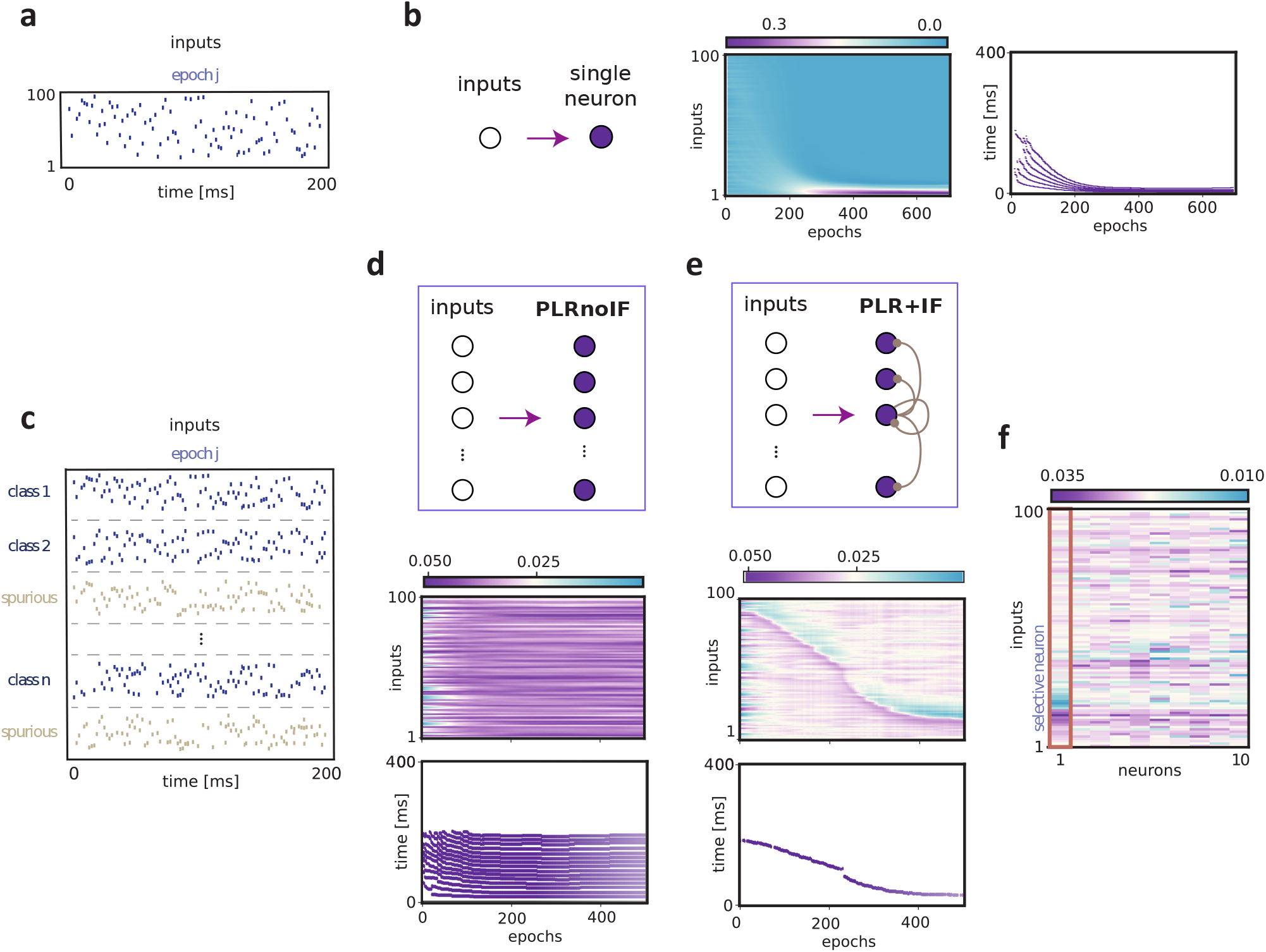
Anticipation of multiple sequences in a neural network with inhibitory feedback. **a**) The input was composed of one spike sequence, given by the correlated firing of *N*=100 pre-synaptic neurons. The *N* pre-synaptic neurons fired sequentially with relative delays of 2 ms, resulting in a total sequence length of 200 ms. **b**) Left: Illustration of the connections between pre-synaptic inputs and the post-synaptic neuron. Middle: Dynamics of the synaptic weights associated with each pre-synaptic input as a function of training epochs. The synaptic weights are ordered along the y-axis from 1 to 100 following the temporal order of the sequence. Right: Dynamics of the neuron spiking activity as a function of training epochs, as in panel b but for 100 pre-synaptic inputs. **c**) The input was composed of sequences belonging to 30 different classes (*n*_class_ = 30) and of *n*_random_ random spiking sequences. **d**) Top-Left: illustration of the PLRnoIF network. Here, we illustrated 5 neurons and all the connections from the neuron in the center as an example. The network was composed of *N*_*nn*_ = 10 neurons. Each neuron receives feedforward inputs and is not connected to the other neurons. Bottom-Left: Dynamics of the synaptic weights 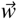 for a neuron in the PLnoGI network (neuron 1) in function of the training epochs, and dynamics of the spiking activity of the same neuron.The synaptic weights are ordered along the y-axis from 1 to 100 following the temporal order of a sequence class (class *c* = 1). PLR+IF: Each neuron inhibits all the neurons in the network via recurrent inhibition. **e**) Same as in **d** for the PLR+IF network. Each neuron receives feedforward inputs and is connected to the other neurons through recurrent inhibition. **d**) The synaptic weight matrix of the PLR+IF network at the end of training (epoch 500). The synaptic weights are ordered along the y-axis from 1 to 100 following the temporal order of the input spikes in sequence class *c* = 1. We highlighted the neuron that developed selectivity to class *c* = 1 (neuron 1), thereby assigning credit to the first inputs of the specific class.

According to the predictive learning rule described above [10], synaptic weights undergo potentiation or depotentiation based on the predictability of the pre-synaptic inputs. However, in cortical networks, neurons may receive many different input sequences. As a result, one would expect that neurons do not selectively fire at the beginning of one particular sequence, but rather diffusely throughout multiple sequences. This raises the question, of how the predictive learning rule can be integrated with competition mechanisms to enable neuronal selectivity for specific spike sequences.

We simulated a case where neurons received input patterns consisting of *n*_class_ distinct sequence classes (Figure 1c). In each sequence, the *N*_presyn_ pre-synaptic neurons fired in sequential order with fixed delays. The total sequence duration was 200 ms. The pre-synaptic neurons participated in each of the *n*_class_ input sequences, and each sequence class *c* ∈ { 1, …, *n*_class_ }was characterized by a unique order of pre-synaptic input spikes. During each training epoch, the input also included randomly drawn sequences whose firing times varied at each iteration, representing irrelevant input patterns (these random sequences comprised 10% of all sequences). We compared two network configurations: In the first configuration, there were *N*_nn_ neurons, but without recurrent connections (“PLRnoIF”). In the second configuration, the neurons were recurrently connected through inhibitory feedback (“PLR+IF”) (see Methods). Specifically, each neuron provided inhibition onto all the neurons in the network, which represents a simplified implementation of the lateral global inhibition in cortical networks [12] (see Methods). In both network configurations, the weight updates followed the predictive learning rule [10]. Furthermore, the synaptic connections from the pre-synaptic inputs were randomly assigned at the start of training, while the weights of the recurrent connections remained fixed.

We exposed the neural network to the input patterns, and we examined the network activity across epochs. In the absence of inhibitory feedback (PLRnoIF network), the neurons showed prolonged firing during the whole duration of each sequence (Figure 1d). The dynamics of the synaptic weights did not show any structure, as all the pre-synaptic inputs became potentiated, which is due to the fact that many sequences were presented. By contrast, we observed that including inhibitory feedback (PLR+IF network) between neurons led to the anticipation of input sequences (Figure 1e). Specifically, neurons in the PLR+IF network became selective to one sequence class and learned to potentiate the early inputs of that sequence (Figure 1f). Accordingly, the neurons in the PLR+IF network fired shortly after the onset of the selected sequence (Figure 1c), and typically did not fire when exposed to the other input sequences (Figure S2). The percentage of neurons that became selective to specific sequences increased across epochs (Figure 2a). By contrast, in the PLRnoIF, no sequence selectivity was developed. As a result of the sequence selectivity, the average firing activity of the neurons in the PLR+IF network decreased substantially (Figure 2b). Importantly, the average duration of the network activity decreased, as many neurons learned to fire at the beginning of distinct sequences (Figure 2c). Additional analyses showed that the network was able to develop sequence selectivity for a vast range of weight initialization (Figure S3a), for different numbers of random sequences in the input pattern (Figure S3b), and for multiple sets of model parameters (Figure S3c).

**Figure 2:**
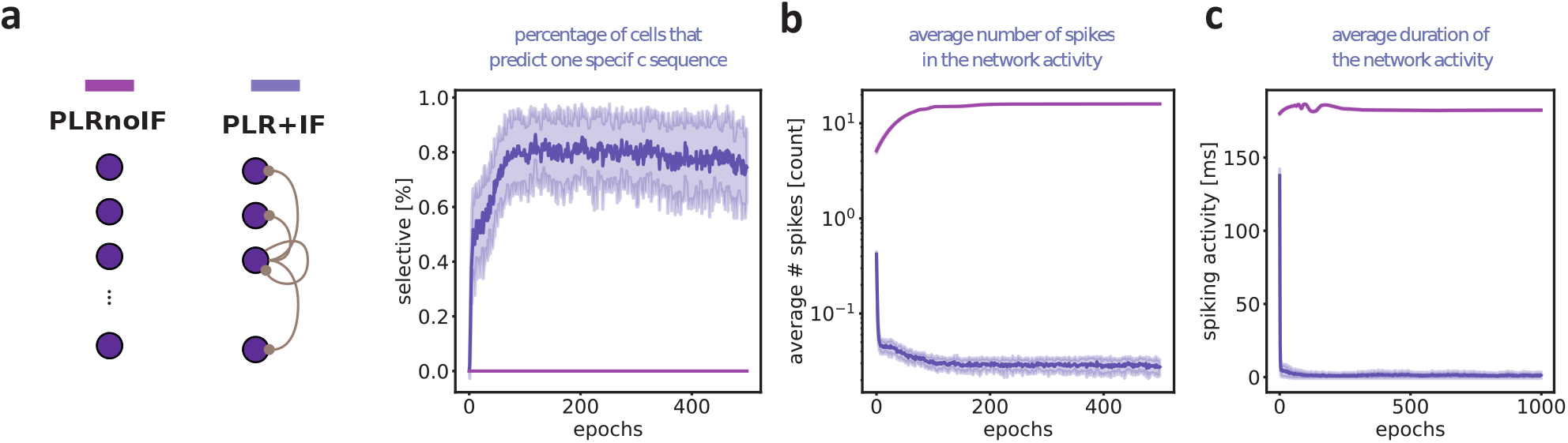
Inhibitory feedback support network selectivity and efficient firing. **a**) Left: Illustration of the two networks and color code. Right: Percentage of selective neurons in the network as a function of the training epochs. A neuron in the network is labeled as *selective* if it fires only for one specific class. The panel shows the percentage of selective neurons for the PLRnoIF and PLR+IF networks, see the color code on the left. **b** The average number of spikes in the network activity as a function of the training epochs. The total number of spikes is averaged across neurons in the network and across sequence classes (*n*_class_ = 30). **c** The average duration of network activity as a function of the training epochs. We computed the temporal difference between the last spike and the first spike of the network activity to estimate the total duration of the network’s activity. The duration of network activity is averaged across neurons and across sequence classes, as in the middle plot. Each panel shows the mean and standard deviation computed over 100 different simulations, and have the same color code as in panel **a**.

Together, our results demonstrate that combining the predictive learning rule with inhibitory feedback leads neurons to fire at the start of particular sequences in the input spike train. The increased selectivity is accompanied by the efficient encoding of input sequences.

### Intermediate levels of inhibition maximize network selectivity and sequence anticipation

In the previous section, we studied the effect of inhibitory feedback in a network of neurons that receives a feedforward input consisting of many different spike sequences. In the PL+GI network, the neurons developed selectivity for specific spike sequences, leading to the anticipation of the selected sequences [10]. These results likely depend on the specific type of interaction between the neurons, in particular on the overall strength of inhibitory feedback. Thus, our next question was how the level of inhibition influences the development of sequence selectivity.

We conducted simulations similar to those in the previous section and systematically varied the strength of inhibitory feedback (Fig 3a). We then examined the percentage of neurons selective to a single sequence at the end of training. The percentage of selective neurons showed an inverted U-curve dependence on the strength of inhibition. Specifically, we observed that the highest percentage of selective neurons was obtained for intermediate values of the inhibition strength, whereas weak or strong inhibition resulted in minimal network selectivity (Fig 3b). The average number of spikes and the duration of network activity decreased progressively as the inhibition strength increased (Fig 3c). The network activity was primarily concentrated at the beginning of each sequence within the same range of inhibition strength (Fig 3d).

**Figure 3:**
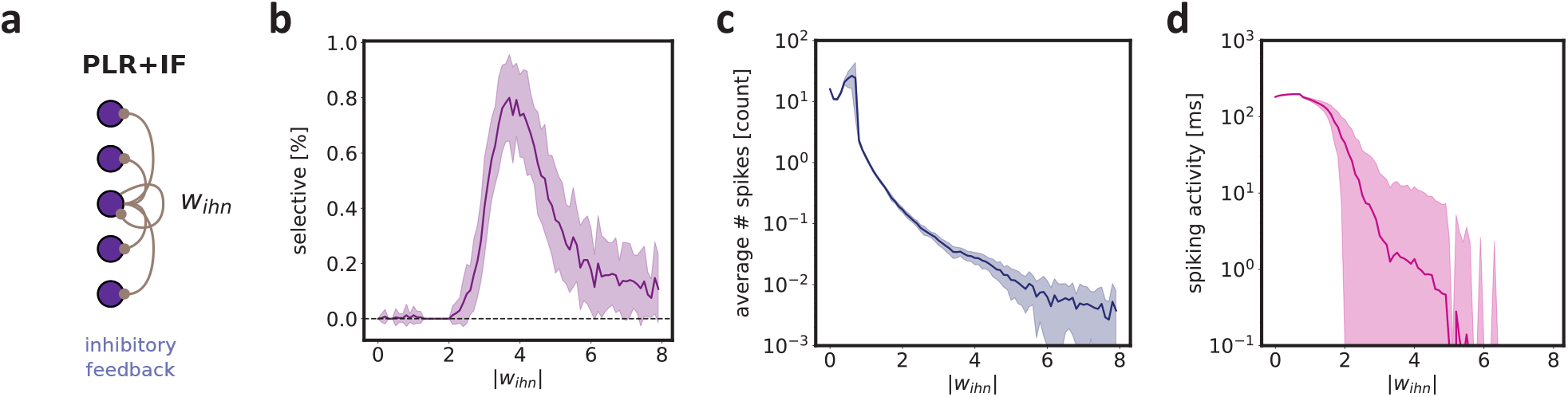
Selectivity and anticipation depend on the strength of inhibition. **a**) Illustration of the global inhibition. The parameter *w*_*ihn*_ defines the strength of the recurrent connections (see Methods). **b**) Percentage of selective neurons as a function of the inhibition strength. A neuron is labeled as *selective* if it fires only for sequences belonging to one specific class. **c**) The average number of spikes in the network activity as a function of the inhibition strength. The total number of spikes is averaged across neurons in the network and across sequence classes. **d**) The average duration of network activity as a function of the inhibition strength. The duration of network activity is averaged across neurons and across sequence classes. Each panel shows the mean and standard error of the mean computed over 1000 different simulations.

Thus, we found that without inhibitory feedback, neurons fired for every sequence, while excessive inhibitory feedback caused the network activity to collapse. However, we observed that an intermediate level of inhibition was optimal for achieving maximum decorrelation among neurons and enabling the anticipation of predictable inputs. Our results suggest that precisely tuned inhibition is a powerful mechanism to maximize the capacity of neural networks.

### Inhibition leads to sequence classification with decorrelated and sparse network activity

We found that when combined with a predictive learning rule, inhibitory feedback promotes sparse activity in the network, and allows individual neurons to fire only for specific sequences. This finding would suggest that a decoder should be able to read out the activity of the recurrent network of neurons and correctly classify the spiking sequences.

To address this, we considered a neural network consisting of two modules: (1) An intermediate PLRnoIF or a PLR+IF layer, and (2) an additional readout layer, where each neuron was assigned to a specific sequence class in the input (Figure 4a). The weights from the intermediate layer to the readout layer were trained using the Adam optimizer with a cross-entropy objective function on the membrane potentials of the readout neurons (see Methods). We exposed the network to sequences belonging to one of ten different classes and evaluated its performance on a dataset where half of the examples did not belong to any specific sequence class (Figure 4b). The input was composed of sequences belonging to *n*_class_ = 10 different classes, and of *n*_random_ random spiking sequences. We used a training dataset with *n*_random_/*n*_class_ = 1 and a testing dataset with *n*_random_/*n*_class_ = 10.

**Figure 4:**
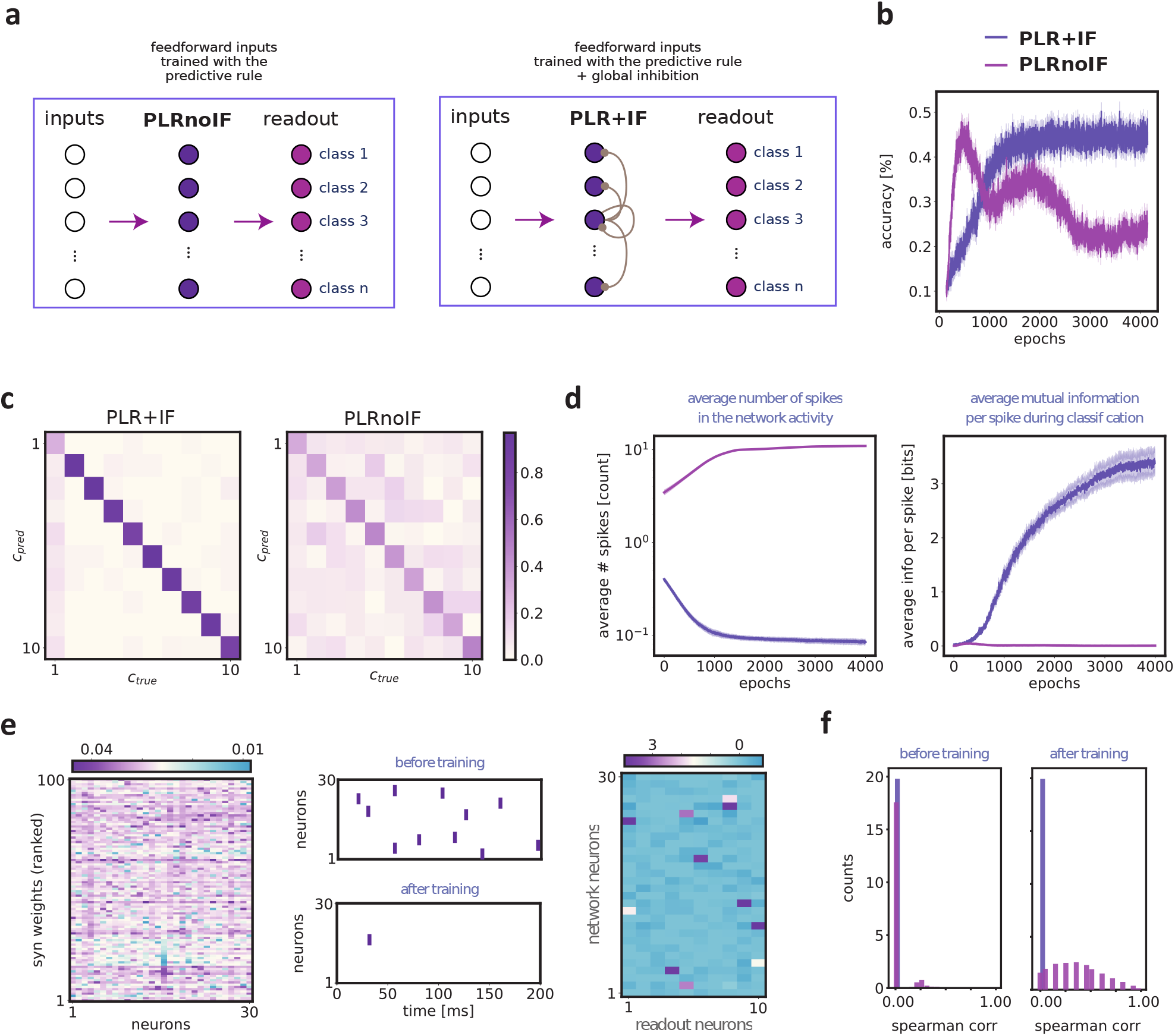
Classification of spike sequences with the predictive learning rule. **a**) The system was composed of an intermediate network (either a PLRnoIF or a PLR+IF network) with *N*_*nn*_ = 30 neurons and a readout layer with *N*_*out*_ = 10 neurons. We updated the synaptic connections from the input to the network following the predictive learning rule, while we updated the connections from the network to the readout layer with backpropagation-through-time (BPTT), see Methods. **b**) Classification accuracy of the readout layer. The panel shows the mean and standard error of the mean computed over 100 different simulations. **c**) Confusion matrix at the end of training. **d**) Left: The average number of spikes in the intermediate network activity as a function of the training epochs. The total number of spikes is averaged across neurons in the network and across sequence classes (*n*_class_ = 10). Right: The average mutual information per spike in the classification task. The mutual information is computed with the conditional probability obtained from the confusion matrix (see Methods). All plots show the mean and standard error of the mean computed over 100 different simulations and have the same color code as in **b. e**) Left) Synaptic weight matrix of the PLR+IF intermediate network after training (epoch 10000) in one example trial. The y-axis represents the ordering of synaptic weights from 1 to 100, following the temporal order of sequence class *c* = 9. The neuron *n* = 16 exhibits selectivity for sequence class *c* = 9, assigning credit to its first inputs. Center) Network activity associated with sequence class *c* = 9 before training (first epoch) and after training (epoch 10000). Right) The synaptic weight matrix of the readout layer at the end of training (epoch 10000). **f**) Distribution of pairwise Spearman correlation coefficients between spike trains of the intermediate network, respectively at the beginning of training (left, epoch 0) and at the end of training (right, epoch 4000). The correlation coefficients are computed from spike trains obtained with 100 different simulations. The color code is as in **b**.

For the PLRnoIF network, we first observed an increase in the classification accuracy and later a decay in performance during training (Figure 4b). This decrease in performance indicates a loss of information in the recurrent network about the sequence inputs. Conversely, the PLR+IF network showed a stable increase in the classification accuracy which was greater than for the PLRnoIF network (Figure 4b-c). This increase in classification accuracy was accompanied by a rise in the percentage of selective cells in the network (Figure S4a). The classification accuracy was driven by a simultaneous decrease in both the prediction error in the intermediate network and the readout layer’s classification error, during both the training and the testing phase (Figure S4b). The average firing activity was two orders of magnitude lower in the PLR+IF network, such the average amount of information per spike was considerably higher than in the PLRnoIF case (Fig 4d).

We further illustrated how synaptic plasticity in both the intermediate network and the readout layer leads to fast sequence classification (Figure 4f). We found that in the PLR+IF network neurons fired in response to the initial inputs of specific sequences, allowing for early classification of the sequence identity (Figure 4f). As a result, only a few early spikes from a small group of neurons were sufficient to achieve maximal classification accuracy, resulting in a sparse connectivity matrix at the readout (4f).

The observation that the PLR+IF network had substantially higher classification performance despite much sparser activity suggests that the input was encoded with a much higher degree of statistical independence between neurons. Previous work has indeed suggested that inhibitory feedback leads to a strong reduction in firing rate correlations among excitatory neurons [16]. We, therefore, quantified the pairwise (Spearman) correlations between all the neurons in the intermediate network module. Firing-rate correlations were much stronger in the PLRnoIF network, and were close to zero in the PLR+IF network (Fig 4e). Hence, inhibitory feedback led to the maximization of channel capacity and efficient encoding of the sequential inputs.

In sum, these analyses show that the predictive learning rule combined with inhibitory feedback leads to improved classification of sequence inputs as compared to a network without inhibitory feedback. Furthermore, predictive learning with inhibition leads to sparse coding with a high degree of information per spike and decorrelated firing between neurons.

### Fast and efficient classification during predictive plasticity

Finally, we compared the (unsupervised) predictive learning rule with two different supervised learning algorithms: BPTT and BPTT with fixed inhibitory connections (BPTT+IF). In the BPTT network, all synaptic connections in both the intermediate network module and the readout layer were trained based on a global loss (cross-entropy) via backpropagation-through-time [17] (Figure 5a). In the BPTT+IF network, the recurrent network module had a fixed level of global inhibition (as in the PLR+IF network, Figure 5a).

**Figure 5:**
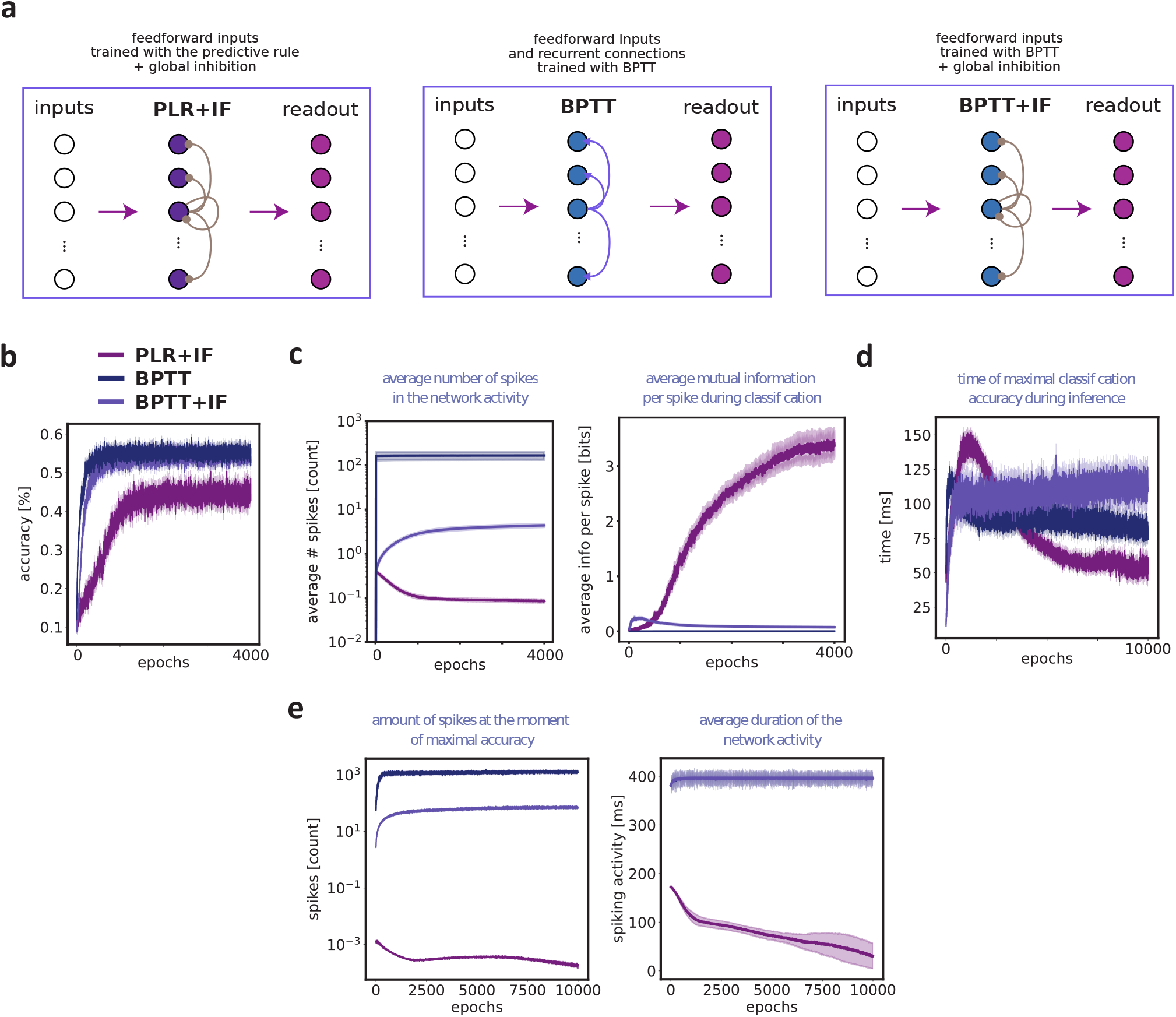
Comparison of the predictive learning rule with back-propagation-through-time algorithms. **a**) Illustration of the three systems. PLR+IF: the system was composed of an intermediate PLR+IF network and a readout layer. We updated the synaptic connections from the input to the network following the predictive learning rule, while we updated the connections from the network to the readout layer with backpropagation-through-time. BPTT: the system was composed of an intermediate network with all-to-all recurrent connections and a readout layer. We updated the synaptic connections from the input to the network, the recurrent connections in the intermediate layer, and the connections from the network to the readout layer with backpropagation-through-time. BPTT+GI: the system was composed of an intermediate network with recurrent connections as in the PLR+IF network. We updated the synaptic connections from the input to the network and the connections from the network to the readout layer with backpropagation-through-time. As in the PLR+IF network, the recurrent connections in the intermediate layer were kept fixed. **b**) Classification accuracy of the readout layer across training. The legend indicates the different training procedures employed for the network. The panel shows the mean and standard deviation computed over 100 different simulations. **c** Left: The average number of spikes in the network activity as a function of the training epochs. The total number of spikes is averaged across neurons in the network and across sequence classes (*n*_*class*_ = 10). Right: The average mutual information per spike in the classification task. The mutual information is computed with the conditional probability obtained from the confusion matrix (see Methods). **d**) The average time step at which the classification accuracy reached its maximum value. **e**) Left: The average number of spikes observed by the readout layer prior to reaching the maximum accuracy. Right: The average duration of network activity. In panel **c, d**, and **e**, all plots show the mean and standard error of the mean computed over 100 different simulations, and have the same color code as in **b**.

We found that the BPTT and the BPTT+IF networks reached a higher classification performance than the PLR+IF network (about 10% higher) (Figure 5b), which is expected considering that both networks were trained using a supervised learning rule. Nevertheless, the PLR+IF network exhibited several features that were absent in the other networks. First, the average firing activity was orders of magnitude lower in the PLR+IF network, and the amount of information per spike was considerably higher (Figure 5c). Second, the readout layer consistently attained maximum accuracy at progressively earlier times in the trial (Figure 5d). In other words, the PLR+IF network signaled the sequence information earlier in time as compared to the network trained with backpropagation-through-time. Third, the number of spikes required to achieve maximal accuracy varied significantly among the different networks (Figure 5d, left). In particular, the number of spikes in the PLR+IF network rule was about 4 orders of magnitude lower than in the other two networks. This result even held true when we analyzed a network trained with BPTT that included regularization (Figure S5). Finally, the duration of network activity was substantially lower in the PLR+IF network (Figure 5d, right).

## Discussion

In this study, we investigated how ensembles of neurons can selectively and efficiently encode multiple temporal input sequences. We implemented a predictive learning rule that adjusts synaptic strengths based on the predictability of pre-synaptic inputs [10]. The main contribution of the present work is to combine the predictive learning rule with recurrent inhibition in a network of neurons. We show that including recurrent inhibition leads to sparse firing in the network, where neurons selectively fire at the beginning of specific sequences. The selectivity of the network was maximized with an intermediate level of inhibition. Moreover, recurrent inhibition led to decorrelated firing between neurons with a high degree of information per spike. Finally, the network activity could correctly reveal the sequence identity, even with just a few early spikes when compared to end-to-end training. Overall, our results demonstrate that fast and energy-efficient classification of different input components can be achieved through the interplay of self-supervised predictive learning rules and inhibitory feedback. The combination of these two mechanisms leads to distinct network activities, which can be used to classify temporal patterns and anticipate their occurrence.

The two main components of our computational model are the predictive learning rule and global inhibition. Neurons endowed with this learning rule learn to fire for the initial, unpredicted inputs in a spiking sequence, resulting in an efficient representation with reduced post-synaptic firing. Importantly, this rule also describes various synaptic plasticity mechanisms that are observed experimentally [10]. Indeed, the synaptic updates crucially depend on changes in the post-synaptic membrane potential that occur at different moments in time, which are believed to trigger LTP and LTD in biological neurons [18, 19]. The predictive learning rule can thereby reproduce STDP mechanisms observed *in-vitro* and describe network phenomena that are usually modeled with STDP rules [2, 4]. For example, the standard asymmetrical STDP window and symmetrical (Hebbian) learning windows come about as a result of predictive plasticity. The impact on synaptic plasticity of pairing frequency [20], initial synaptic strength [21] and higher-order spike patterns (beyond pre-post pairing) [22, 23] are also described by the predictive plasticity rule. Importantly, this rule was derived from an optimization problem based on predictive processes, thus suggesting that STDP is a consequence of a general learning rule given the particular state of the system and the stimulation protocol [24]. We refer to [10] for a detailed analysis of the relation between the predictive plasticity rule and STDP models and experiments.

The inclusion of global inhibition in our model is well-supported by experimental evidence, indicating patterns of massive convergence and massive divergence from excitatory neurons to interneurons, e.g. fast-spiking parvalbumin-positive (PV) cells [25], without connection specificity. These interneurons span several cortical columns and provide fast and unspecific inhibition to pyramidal neurons [26]. Our model captures the essence of an excitatory-inhibitory network model, where single neurons adjust their synaptic weights based on local learning rules. At present, our model did not include recurrent excitation, which is a hallmark of cortical networks [27, 28, 29] but is typically absent in subcortical areas. Thus, in the present form, the model presented here may best approximate unsupervised learning in subcortical networks. Future works should include recurrent excitation in the model and investigate its impact on neuronal selectivity and anticipation of spike sequences.

The recurrent inhibition implemented in this study resembles the mechanism known as *k*-Winner-Take-All (*k*-WTA) with *k* = 1, where only one neuron emerges as the winner, and represents a specific pattern [11]. Experimental evidence demonstrates that lateral inhibition in cortical networks enables this type of operation [30] while also promoting decorrelation of neuronal activity [16]. In agreement with experimental and theoretical findings [31, 32, 16], we found that global inhibition decreases the correlation between neurons, and provides a mechanism for independent encoding of multiple spike sequences. Our results also indicated that an intermediate level of inhibition is optimal for decorrelated firing in the network with high information content per spike. This suggests that there might be a vulnerability of functional cortical activity to GABAergic disturbances or excitation-inhibition balance [33]. Indeed, experimental findings showed that extreme levels of inhibition are associated with psychiatric diseases and that intermediate inhibitory strength correlates to functional brain activity [34, 33].

However, in cortical networks, different cell types contribute to lateral inhibition, exhibiting diverse neuronal properties and connectivity patterns with excitatory neurons [35, 36, 37]. This heterogeneity gives rise to various excitation-inhibition motifs [38, 39], transient and oscillatory activity [40, 41], and significantly affects synaptic plasticity [42]. Therefore, it is important to note that our current work employs a simplified implementation of recurrent inhibition in the network. Future works will incorporate inhibitory neurons into the network and explore the role of different connectivity schemes in anticipating multiple sequences. This will also provide insights into how the prediction of multiple sequences relates to the balance of excitation and inhibition in neural networks. Furthermore, the emerging evidence of inhibitory plasticity [43, 44] and its diverse consequences on cortical computations [45] introduces another interesting direction for exploration. It is e.g. possible that inhibitory plasticity contributes to reaching an optimum, i.e. intermediate level of global inhibition.

Previous studies have investigated the combination of local learning rules and competition mechanisms. For instance, certain forms of spike-timing-dependent plasticity (STDP), when combined with global winnertake-all mechanisms, can result in neurons representing different segments of a repeating pattern [13, 46]. Additionally, specific STDP windows can produce a reduction in output latency for frequently occurring inputs [8, 47]. Other spike-based learning rules, when combined with WTA mechanisms, have also shown high performance in classification tasks [48, 49]. It is worth noting that the predictive learning rule encompasses multiple STDP phenomena, including multiple STDP windows and higher-order effects (beyond pre-post pairing) [10]. Thereby, our findings provide a general description of sequence anticipation as a result of predictive processes based on synaptic plasticity, whereby STDP mechanisms come about as a consequence of predictive plasticity. Differently from previous works, we also performed a direct comparison of the predictive learning rule with standard training algorithms, and we showed that the combination of self-supervised learning rules and global inhibition led to fast, and energy-efficient encoding of stimulus identity compared to backpropagation-through-time (BPTT).

Our model relies on the temporal relationships between input spikes, and it has been tested on sequential pre-synaptic spike trains. Experimental evidence from various brain regions supports the existence of reliable spike sequences [3, 50]. For instance, the serial firing of neurons can represent the sequential nature of external stimuli. Moving objects can trigger the sequential firing of neurons in the visual system [4], resulting in waves of activity that can contain predictive information about the object’s trajectory [51, 52]. In the hippocampus, the sequential activity of place cells encodes the motion of the animal entering different locations [53]. Moreover, spike sequences can also emerge as internally generated activities, e.g. during goal-oriented behaviour [54]. Neurons can engage in structured sequential firing during internal simulations and memory consolidation [55], motor planning, and online control of actions [54]. Interestingly, these sequential activities exhibit experience-dependent facilitation [2], and can be replayed at compressed time scales [56, 5]. Moreover, reliable spike sequences can be induced only by the activation of a few spikes neurons [57, 58]. Our findings support the idea that even a few spikes can initiate a cascade of neuronal activity and that learning can flatten internal activity by assigning credit to inputs with predictive power. Thereby, our model provides insight into the neuronal mechanisms underlying reliable spike sequences and their experience-dependent compression at the network level. Our results suggest a functional role of sequential spiking activity for predictive processes and for the engagement of networks in anticipating future events and organizing behaviour [2].

Finally, the results presented in this study have implications for training spiking neural networks (SNNs) with biologically-inspired algorithms, particularly for assigning credit to local synapses based on non-local information [59]. Indeed, the credit-assignment problem is particularly hard in SNNs, where the errors are propagated both in time and in space and through non-differentiable activation functions [60, 61]. Previous research has approached this problem by drawing insights from neuroscience, including the interplay between different neuronal types [62, 63], the multiplexing of signals through distinct firing patterns [63] and the involvement of different neuronal compartments [64, 65, 66]. Here, we showed that the combination of a local, self-supervised learning rule and an arguably simple inhibitory mechanism leads to distinguishable network activities and successful classification of neuronal inputs. Moreover, the predictive learning rule enables gradient propagation through differentiable functions. Indeed, the loss functions at the single neuron level depend solely on the membrane potential of each respective cell. It is important to note that our model was specifically tested on classifying spike sequences with direct temporal relationships between input spikes. An interesting direction for future research would be to apply our model to different classification tasks and systematically evaluate its performance in various scenarios.

## Methods

### Network model

We implemented a recurrent neural network composed by *N*_*nn*_ neurons, where each *i*-th neuron in the network received external inputs from *N*_presyn_ pre-synaptic neurons 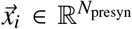. The synaptic weights associated with the recurrent connection and with the external inputs were determined by the connectivity matrix 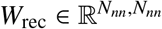 and 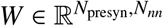, respectively. Each *i*-th neuron in the network obeyed a discrete-time model of the form

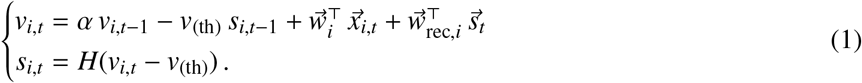

Here, *v*_*i,t*_ ∈ ℝ is the membrane potential of the *i*-th neuron at timestep *t*, α ≡ 1 −*h*/τ_*m*_ where τ_*m*_ is the membrane time constant and *h* is the timestep size, 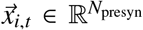 is the pre-synaptic input to the *i*-th neuron at timestep *t, v*_(th)_ is the spiking threshold (the subscript _(th)_: “threshold”), and *H*(·) is the Heaviside function. The weight vector of the external inputs 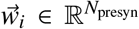 and of the recurrent connections 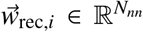are the *i*-th column of the connectivity matrix *W* and *W*_*rec*_, respectively. The two connectivity matrices *W* and *W*_*rec*_ were defined depending on the specific type of simulations, see the section *Optimization and training schemes*. The variable *s*_*i,t*_ ∈ {0, 1 }takes binary values and indicates the presence or absence of an output spike at timestep *t*. If the voltage exceeds the threshold, an output spike is emitted, and this event reduces the membrane potential by a constant value *v*_(th)_ at the next time step. This implementation of the membrane potential reset relates our model to the spike response model [67]. We set *h* = 1 ms in all numerical simulations.

We also implemented a readout layer composed of *n*_class_ neurons, one for each class *c*. Each *i*-th neuron in the layer received the activity of the recurrent neural network 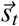 and the associated synaptic weights were determined by the connectivity matrix 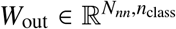. The readout neurons obeyed a discrete-time equation of the form

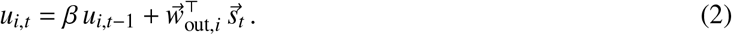

Here, β ≡ 1 − *h*/τ_*m,out*_ where τ_*m,out*_ is the membrane time constant of the readout neurons and *h* is the timestep size, and 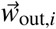 is the *i*-th column of the connectivity matrix *W*_out_.

### Predictive learning rule

The predictive learning rule used in this study was adapted from a previously published paper [10]. This rule is derived from an optimization problem in time at the single neuron level. Specifically, each *i*-th neuron in the network is assigned with an objective function ℒ_*i*_ as follows,

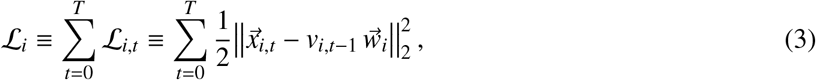

where || · ||_2_ is the *l*_2_-norm, 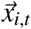_*t*_ are the external inputs to the *i*-th neuron in the network, *v*_*i,t*−1_ is the membrane potential of the *i*-th neuron and 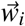is the synaptic weight vector of the external inputs, see Equation (1). Here, the objective is to obtain the minimal difference between the input 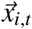 received by the neuron, and its prediction via *v*_*i,t*−1_ and 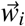. To derive the predictive learning rule, one computes the gradient of ℒ_*i*_ w.r.t. 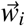,

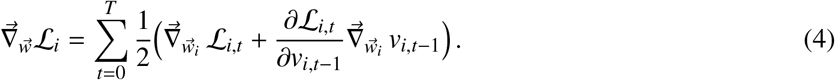

The first term accounts for the direct effect of a weight change on ℒ_*t*_, while the second accounts for its indirect effect via the membrane potential *v*_*i,t*−1_. The first term of the gradient is given by

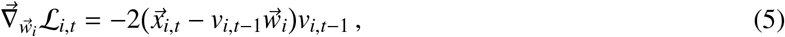

while the second term is given by

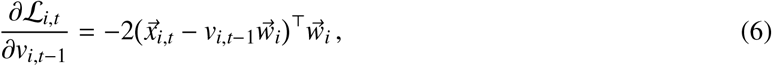

and by

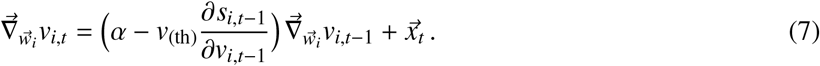

We define an influence vector 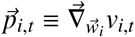 such that it obeys the recursive equation (see Equation (7))

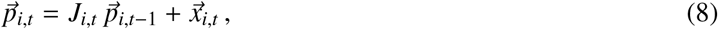

where *J*_*i,t*_ is the Jacobian from the recurrent equation of the membrane voltage of the *i*-th neuron in Equation (1)

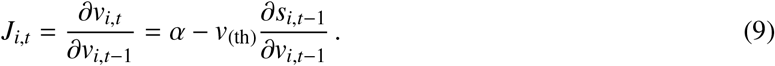

The gradient of ℒ_*i*_ w.r.t. 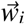is then given by

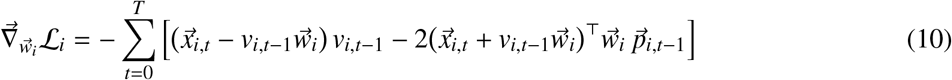

We define the prediction error 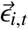at timestep *t* as

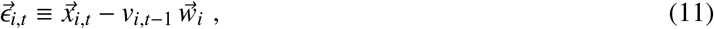

that determines the sign and amplitude of synaptic plasticity, and the global signal ε_*i,t*_ at timestep *t* as

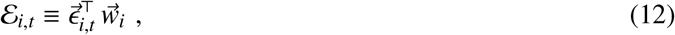

thereby the gradient of ℒ_*i*_ w.r.t. 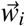is given

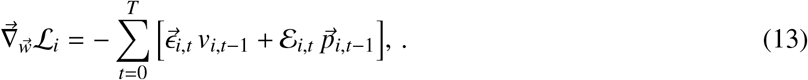

After the end of the period [0, *T*], the exact gradient can be used to update the weight vector via gradient descent.

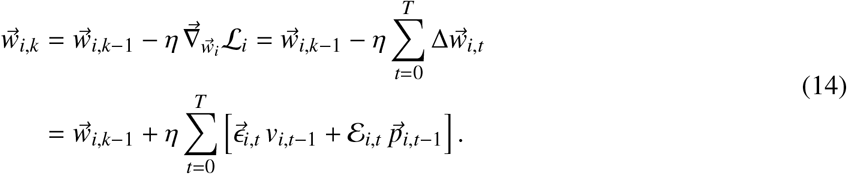

where η is the learning rate, and 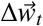is the weight update obtained with the predictive learning rule [10]

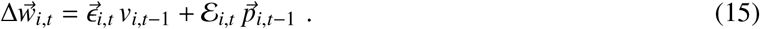

Each neuron in the network receives inputs from other neurons, and the associated recurrent connections are defined by the specific connectivity scheme. The contributions to the gradient of the recurrent connections from neuron *j* to neuron *i* in the network are proportional to

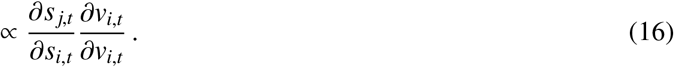

These contributions have a discontinuous effect at the time of output spikes, and thereby we neglected these contributions to the gradient. Conforming to previous works [68, 61, 69], we also neglected the contribution of the reset mechanisms in Equation (9), that is

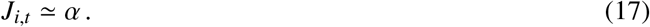

In the predictive learning rule, the membrane potential *v*_*i,t*_ serves as the input variable for the loss function, while *s*_*i,t*_ acts as a hidden variable. Thus, the predictive learning rule circumvents the issue of back-propagating through discrete output variables.

In this work, we also considered weight updates that take place in real time with the prediction of the presynaptic inputs. To do so, we approximated the Equation (14) with the current estimate of the gradient at each timestep *t*,

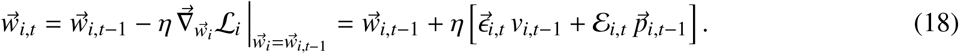

### Spike sequences datasets

Each dataset is composed of spike sequences belonging to *n*_class_ different sequence classes, and comprise the correlated activity of *N*_presyn_ pre-synaptic neurons. Each sequence class *c* ∈ *n*_class_ is defined by a unique order of the firing times of the pre-synaptic neurons. The *N*_presyn_ pre-synaptic neurons fire one spike in each sequence with relative delays of 2 ms, resulting in a total sequence length of 2 *N*_presyn_ ms. We also add to the dataset *n*_random_ random spiking sequences, that is, sequences whose order of firing was drawn randomly. We created the dataset as follows. First, we set the number of pre-synaptic neurons *N*_presyn_ and the number of sequence classes *n*_class_, then we randomly drew the order of firing of the *N*_presyn_ pre-synaptic neuron in each class *c*. Second, we created *b*_class_ examples of the spike sequences belonging to each class *c*. Finally, the inputs were convoluted with an exponential kernel with τ_*x*_ = 2 ms to replicate the fast dynamics of post-synaptic currents. Thereby, the dataset was composed by *N*_(dataset)_ = (*b*_class_ * *n*_class_ + *n*_random_) spike sequences. An example set of spike sequences from *N* = 10 pre-synaptic neurons and belonging to *n*_class_ = 4 classes is as follows:

**Table 1:** Example set of spike sequences from *N* = 10 pre-synaptic neurons and belonging to *n*_class_ = 4 classes.

|  | 1 | 2 | 3 | 4 | 5 | 6 | 7 | 8 | 9 | 10 |
| --- | --- | --- | --- | --- | --- | --- | --- | --- | --- | --- |
| $\mathbf{c_1}$ | 10 ms | 18 ms | 2 ms | 4 ms | 12 ms | 16 ms | 8 ms | 0 ms | 20 ms | 14 ms |
| $\mathbf{c_2}$ | 0 ms | 8 ms | 12 ms | 18 ms | 10 ms | 6 ms | 2 ms | 20 ms | 4 ms | 14 ms |
| $\mathbf{c_3}$ | 14 ms | 10 ms | 20 ms | 4 ms | 2 ms | 8 ms | 16 ms | 6 ms | 0 ms | 12 ms |
| $\mathbf{c_4}$ | 4 ms | 2 ms | 0 ms | 12 ms | 20 ms | 18 ms | 6 ms | 16 ms | 14 ms | 10 ms |

In Figure 1a-b and Figure S1 the dataset was composed by one example (*b*_class_ = 1) of a single sequence class (*n*_class_ = 1), and without examples of random spiking sequences (*n*_random_ = 0).

In Figure 1, Figure 2, Figure 3 and Figure S2 the dataset was composed of several examples (*b*_class_ = 500) for each sequence class (*n*_class_ = 30), and by random spiking sequences (*n*_random_ = 500). In Fig S3 we used the same ratio between class examples and random sequences, but with *n*_class_ = 10.

In Figure 4, Figure 5, Figure S4, Figure S5 and Figure S2 the dataset was composed of several examples (*b*_class_ = 50) for each sequence class (*n*_class_ = 10), and by random spiking sequences (*n*_random_ = 500).

### Objective functions and regularizations

We defined the predictive objective function of a network as

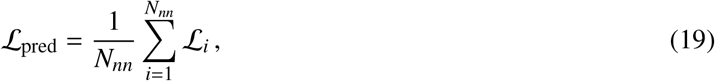

where ℒ _*i*_ is the objective function associated to each *i*-th neuron in the network, see Equation (3).

We trained the readout layer using supervised learning. During each iteration in the training and testing phase, the actual label *y*_*s*_ of every random sequence was drawn randomly from the *n*_class_ possible classes, resulting in the following dataset

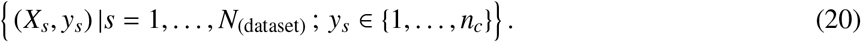

Here, *X*_*s*_ is an example sequence in the dataset and *y*_*s*_ is the associated label. We applied a cross-entropy loss to the activity of the readout layer 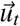,

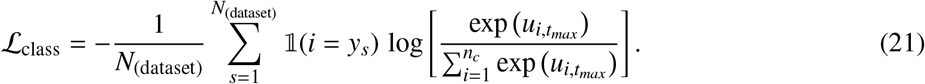

Here 𝟙 is the indicator function and 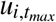 is the maximal membrane potential of the *i*-th readout neuron across timesteps, where *t*_*max*_ = argmax_*t*_ *u*_*i,t*_.

In Figure S5 we also applied an L1-regularization to the total number of spikes emitted by the network, as to penalize excessive firing

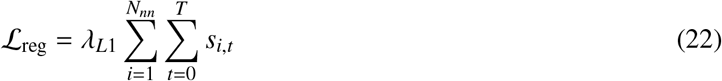

where λ_*L*1_ is a hyperparameter controlling the influence of the regularization term.

### Optimization and training schemes

In the following, we provide a description of the optimization techniques and the training schemes used in this study.

In Figure 1, Figure 3, Figure S1, Figure S2 and Figure S3, we minimized ℒ_pred_ using the online approximation of the predictive learning rule. That is, each *i*-th column of *W* was updated following Equation (18).

In Figure 4, Figure 5, Figure S4, and Figure S5 we used different networks with different optimization schemes. For the case of the predictive neural network, we employed backpropagation-through-time (BPTT) to minimize both the network’s objective ℒ_pred_ and the readout layer’s objective ℒ_class_. For the LIF network with recurrent inhibition, we used BPTT to minimize the objective ℒ_class_. We updated all the synaptic weights in the networks, except for the connectivity matrix *W*_rec_ that was kept fixed. Finally, in the LIF network without recurrent inhibition, we also employed BPTT to minimize the objective ℒ_class_ of the readout layer. Unlike the previous case, we updated all the synaptic weights in the networks.

For all the spiking neural networks that were not optimized with the predictive learning rule, we applied a surrogate gradient to replace the spiking non-linearity with a differentiable function [61]. Specifically, we chose a fast sigmoid function of the form

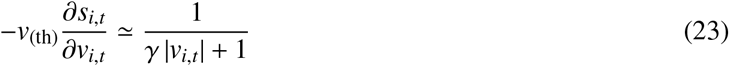

where γ is a parameter controlling the steepness of the sigmoid function. We optimized the model parameters with the Adam optimizer [70] in all the simulations shown in this study.

### Weights initialization

In Figure 1, Figure 2, Figure 3, and Figure S1 the synaptic weights from the input dataset to the network were determined by the connectivity matrix 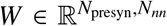. Before training, the entries of *W* were independently drawn from a uniform distribution 𝒰 (0, *w*_*init*_), with *w*_*init*_ > 0. The connections between neurons in the network were defined by the connectivity matrix 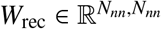. Before training, the connectivity matrix *W* was defined as 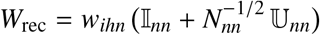, where 𝕀_*nn*_ is the identity matrix, *N*_*nn*_ is the number of neurons in the network, 𝕌_*nn*_ is a random matrix drawn from a uniform distribution between [0,1], and *w*_*ihn*_ < 0 defines the overall strength of the recurrent inhibition. The connectivity matrix *W*_rec_ was fixed during the simulations, that is, the associated synaptic weights were not subject to plasticity.

In Figure 4, Figure 5, Figure S4, and Figure S5 we used different types of networks with different initialization schemes. The connections between the network and the readout layer were determined by the connectivity matrix 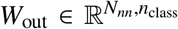. Before training, the entries of *W*_*out*_ were independently drawn from a normal distribution 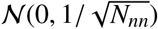. For the case of the predictive neural network and the LIF network with inhibition, we initialized the connectivity matrices *W* and *W*_rec_ as in Figure 1 and Figure 3. For the case of the LIF network without inhibition and the random neural network, the entries of *W* and of *W*_rec_ were independently drawn from the normal distribution 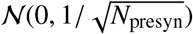and 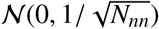, respectively.

### Data analysis

We calculated each entry *C*_*i j*_ of the confusion matrix in Figure 4 as the average number of observations known to belong to sequence class *i* and predicted by the readout to belong to sequence class *j*. We calculated the average entry *C*_*i j*_ at the end of training (epoch 4000) and over 100 different simulations.

To calculate the Mutual Information (MI) in Figure 4, we first assume a uniform prior distribution *p*(*j*) of the sequence class, and we calculated the joint probability *p*(*i, j*) as the product of the prior distribution *p*(*j*) and the conditional distribution *p*(*i*| *j*) obtained from the confusion matrix,

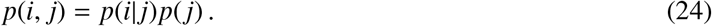

Then, we calculated *p*(*j*) as the sum of the joint probability *p*(*i*| *j*)*p*(*j*) over the expected sequence class *p*(*j*), and we finally calculate the MI as

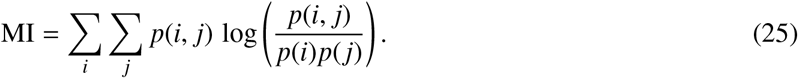

We averaged the MI in each epoch over 100 different simulations.

We calculated the average Spearman correlation corr(*k, l*) in Figure 4 between each pair of spike train *k* and *l* as

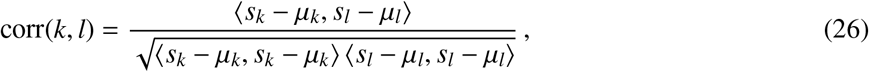

where *s* is the spike train, *μ* is the mean of the respective spike train, and ⟨·⟩ is the dot product. We calculated corr(*k, l*) of the spike trains of any pair of neurons (*k, l*), for each sequence class *c* and for 100 different simulations.

## Acknowledgments

This work was supported by an ERC Starting Grant (SPATEMP) and a BMF Grant (Bundesministerium fuer Bildung und Forschung, Computational Life Sciences, project BINDA, 031L0167). We thank Wolf Singer and Richard Gao for helpful comments, edits, and insightful discussions.

## Authorship contributions

Conceptualization: MS, MV. Mathematical analysis: MS. Simulations: MS. Writing: MS, MV. Supervision: MV.

## Code availability

The code to reproduce all the Figures in the main text and in the Supplementary Materials is freely available at *we will insert GitHub repo link here*.

---

**Figure S1:**
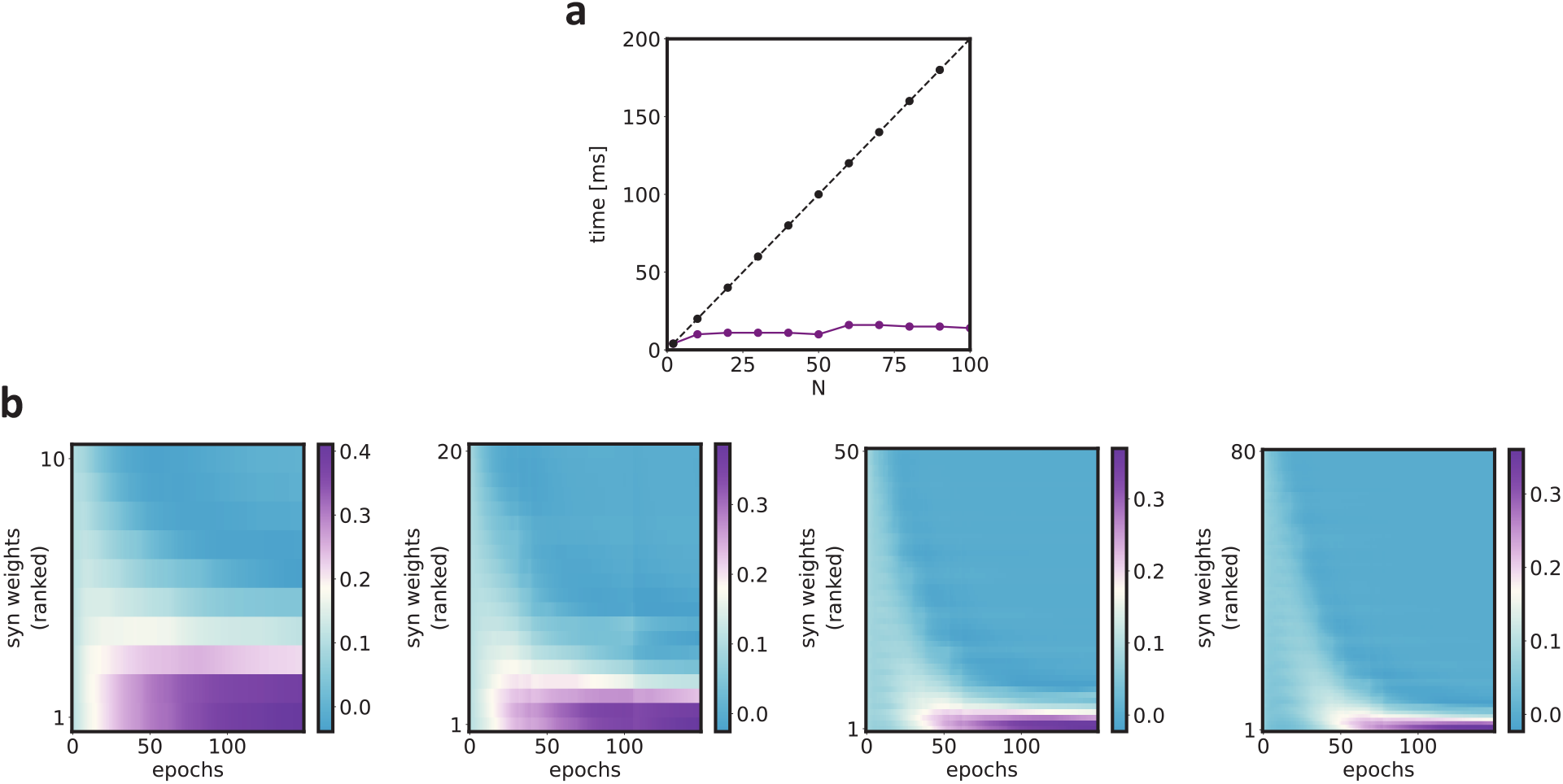
Dependence of sequence anticipation to model parameters - relates to Figure 1 in the main text. **a**) The purple line represents the duration of the neuron activity at the end of training (epoch 150) for different numbers of pre-synaptic neurons *N*. We performed the same simulations with the same model parameters for each value of *N*. The black line represents the duration of the associated pre-synaptic spike sequence. As in Figure 1, the *N* pre-synaptic neurons fired sequentially with relative delays of 2 ms, resulting in a total sequence length of 2* *N* ms in each case. The neuron learns to fire at the beginning of each sequence, regardless of the number of pre-synaptic neurons *N*. **b**) Dynamics of the synaptic weights 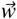 as a function of the training epochs, and for different numbers of pre-synaptic inputs *N*. From left to right: *N* = 10, *N* = 20, *N* = 50, *N* = 80. In each plot, the synaptic weights are ordered along the y-axis from 1 to *N* following the temporal order of the sequence.

**Figure S2:**
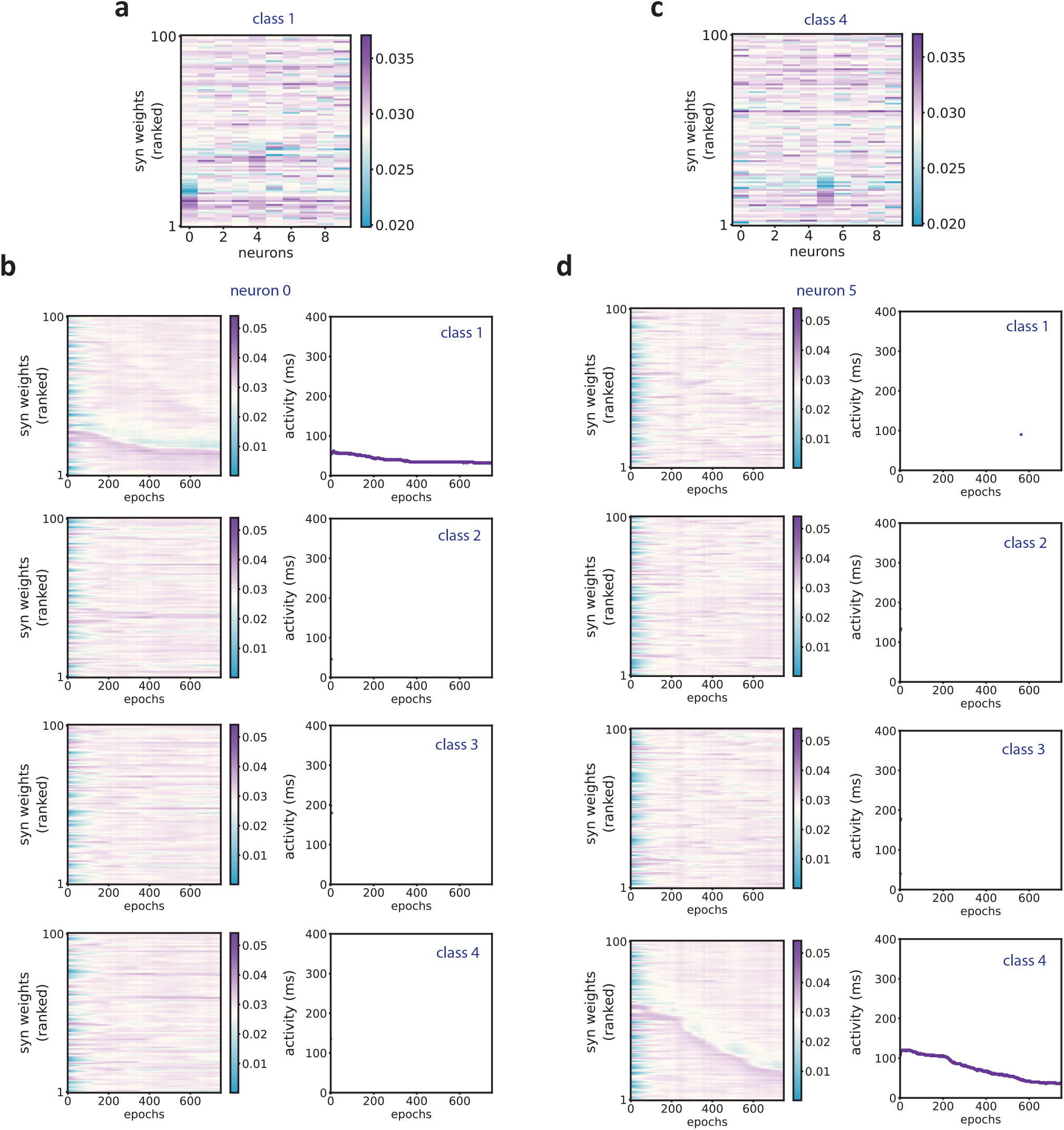
Evolution of the synaptic weights and the spiking activity for selective neurons - relates to Figure 1 in the main text. Simulations were performed as in Figure 1, but with different network initialization. **a**) The synaptic weight matrix of the network at the end of training (epoch 750). The input was composed of sequences belonging to 30 different classes (*n*_class_ = 30). The synaptic weights are ordered along the y-axis from 1 to 100 following the temporal order of the input spikes in sequence class *c* = 1. Neuron *n* = 0 developed selectivity to class *c* = 1, and assigned credit to the first inputs in the order of class *c* = 1. **b** Left) Dynamics of the synaptic weights 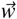 of the neuron *n* = 0 as a function of the training epochs. In each plot, the synaptic weights are ordered along the y-axis from 1 to 100 following the temporal order of the specific sequence class *c*. The synaptic weights dynamics resemble the one in Figure 1d only for class *c* = 1. The neuron developed selectivity due to the recurrent inhibition in the network, thereby assigning credit only to the first inputs of class *c* = 1. Right) Dynamics of the spiking activity of the neuron *n* = 0 as a function of the training epochs. Each plot shows the dynamics of the spiking activity across epochs when the neuron was presented to the specific sequence class *c*. The neuron developed selectivity by firing only when presented to class *c* = 1. The neuron grouped its activity earlier in time, such that it eventually learned to fire for the first inputs in the sequence class *c* = 1. **c**) Same as in **a** for the sequence class *c* = 4. **d**) Same as in **b** for the neuron *n* = 5.

**Figure S3:**
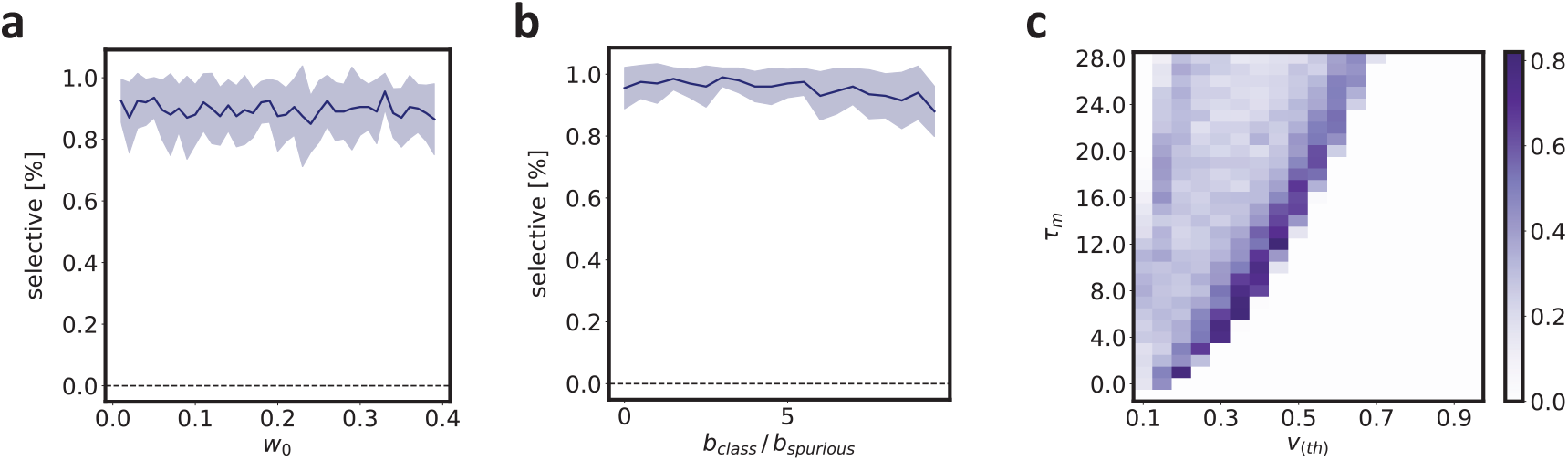
Dependence of network selectivity to model parameters - relates to Figure 1 in the main text. Simulations were performed as in Figure 1, but with *n*_class_ = 10. **a**) Network selectivity is shown as a function of the mean of the initial weight matrix for the external input *W*. The network selectivity does not strongly depend on the initial conditions. For each weight, we performed 100 simulations (shown are the standard errors of the mean). For each simulation, we used 1000 epochs. **b**) As in panel **a**, but as a function of the percentage of random sequences in the dataset. **c**) As in panel **a**, but as a function of the membrane time constant τ_*m*_ and of the spiking threshold *v*_(th)_. Here we performed the simulations with *b*_*class*_/*b*_*random*_ = 1. The network gets selective for a vast range of model parameters. The highest degree of selectivity is obtained at the border of the region where the neurons in the network start spiking.

**Figure S4:**
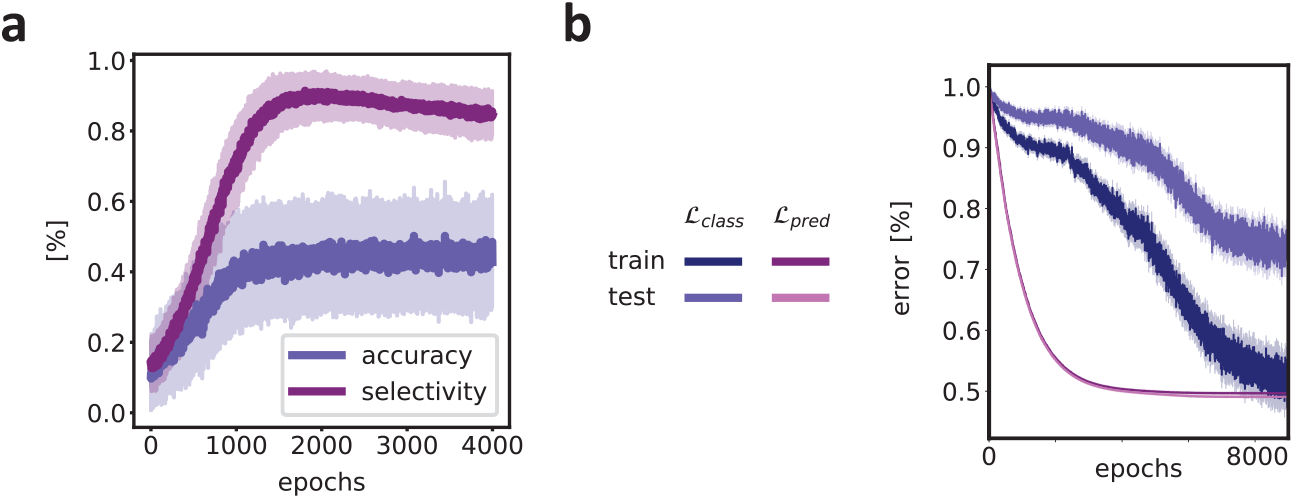
Model performances and network selectivity - relates to Figure 4 in the main text. **a**) Classification accuracy of the readout layer and percentage of selective neurons in the PLR+GI network, see the legend. The panel shows the mean and standard deviation computed over 100 different simulations. **b**) Evolution of the objectives ℒ_class_ andℒ _pred_ during the training and testing phase. The random sequences comprised 10% of the dataset during the training phase, and 50% of the dataset during the testing phase. All the objectives functions are normalized to the initial value at epoch 0.

**Figure S5:**
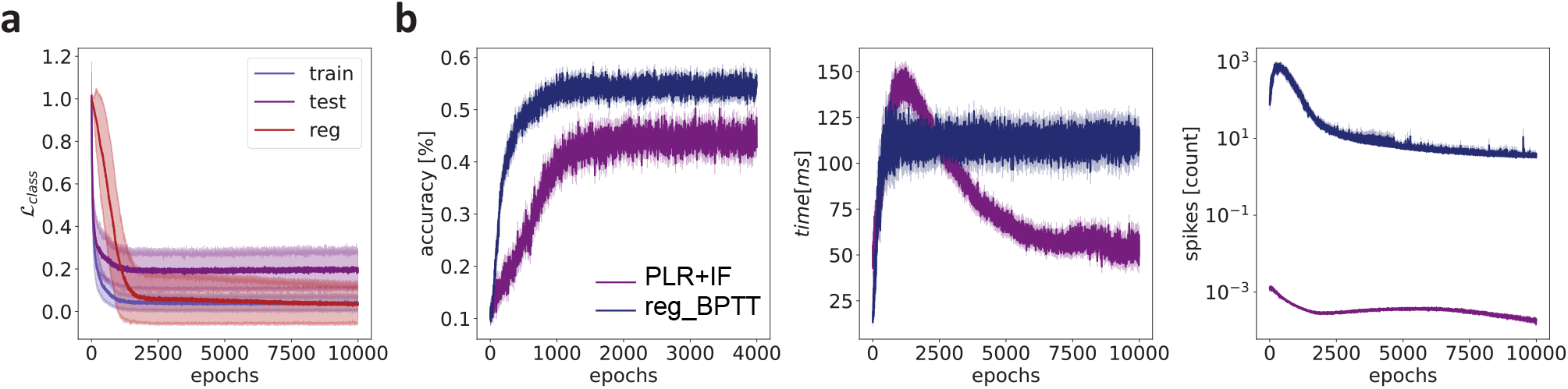
Effect of regularization on the classification of spike sequences - relates to Figure 5 in the main text. Simulations were performed as in Figure 5, but here we compared the performance of the PLR+GI network with the BPTT network augmented with a *L*1 regularization on the network activity (reg BPTT). Specifically, we trained the network end-to-end on the total objective 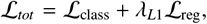, where 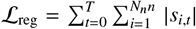 and λ_*L*1_ is a hyperparameter that controls the influence of the regularization term, see Methods. **a**) Evolution of the objectives ℒ_class_ for the train and test sets, and of the objective ℒ_reg_ for the train set. All the objective functions are normalized to the initial value at epoch 0. **b**) The classification accuracy of the readout layer during training is shown on the left. The reg BPTT network achieved similar performances to the BPTT network in Figure 5d. The average time step at which the classification accuracy reached its maximum value is shown in the center. The PLR+GI network reaches its maximal performance in half the time compared to the reg BPTT network. The average number of spikes observed by the readout layer prior to reaching the maximum accuracy is presented on the right. As expected, the regularization term limits the total amount of spikes in the network activity.

